# Designing Biological Micro-Sensors with Chiral Nematic Liquid Crystal Droplets

**DOI:** 10.1101/2021.10.25.465736

**Authors:** Lawrence W. Honaker, Chang Chen, Floris M.H. Dautzenberg, Sylvia Brugman, Siddharth Deshpande

**Affiliations:** Wageningen University & Research, Laboratory of Physical Chemistry and Soft Matter, Stippeneng 4, 6708 WE Wageningen, The Netherlands; Wageningen University & Research, Host–Microbe Interactomics Group, De Elst 1,6708 WD Wageningen, The Netherlands

## Abstract

Biosensing using liquid crystals has a tremendous potential by coupling the high degree of sensitivity of their alignment to their surroundings with clear optical feedback. Many existing set-ups use birefringence of nematic liquid crystals, which severely limits straightforward and frugal implementation into a sensing platform due to the sophisticated optical set-ups required. In this work, we instead utilize chiral nematic liquid crystal micro-droplets, which show strongly reflected structural colour, as sensing platforms for surface active agents. We systematically quantify the optical response of closely related biological amphiphiles and find unique optical signatures for each species. We detect signatures across a wide range of concentrations (from *μ*M to mM), with fast response times (from seconds to minutes). The striking optical response is a function of the adsorption of surfactants in a non-homogeneous manner and the topology of the liquid crystal orientation at the interface requiring a scattering, multidomain structure, which we observe to be different between molecules. We show lab-on-a-chip capability of our method by drying droplets in high-density two-dimensional arrays and simply hydrating the chip to detect dissolved analytes. Finally, we show proof-of-principle *in vivo* biosensing in the intestinal tracts of live zebrafish larvae, demonstrating CLC droplets show a clear and differential optical response between healthy and inflamed tissues. Our unique approach has great potential in developing on-site detection platforms and detecting biological amphiphiles in living organisms.

## Introduction

Widely used in digital display applications^1^, liquid crystals (LCs), ordered fluid phases formed from strongly anisotropic molecules, have become increasingly popular for use in chemical and biological sensing applications^2–9^. Many of the same qualities that have lent themselves to their use in displays – extreme sensitivity to aligning conditions, rapid switching and response times, and a clear optical response – make them well-positioned for their use in sensing applications. Most sensing research to date has used nonchiral nematic LCs (NLCs)^3–6,8^, looking at either the switching between alignment configurations or the changes in the LC phase caused by the adsorption of an amphiphile^2,10,11^ or by the infiltration of a contaminant such as a volatile gas^7,12–14^. Simple prototype sensors have been developed based on the use of the LC either as a primary or as a secondary sensing component^2,15^. These visualize the presence or absence of an antigen through a switch of alignment that becomes reflected in a change in birefringence. A limiting condition is that NLCs function normally as a binary switch, transitioning between a bright “on” state and a dark “off” state. Only in some extreme cases, transient states of tilted alignment can be observed, making intermediate concentrations distinguishable, but a complete change of LC alignment is typically observed at concentrations well below the critical micelle concentration^11,16^: once full switching has occurred, it is difficult to extract further information from an NLC interface.

A less-explored but equally interesting direction is the use of cholesteric liquid crystals (CLCs), also known as chiral nematics, for sensing applications. CLCs, in addition to the alignment-dependent birefringent properties of NLCs, additionally have a helical modulation in their orientation. These are usually formed by mixing a chiral molecule (often called a ‘dopant’) into a nematic phase, with the final equilibrium pitch *p*_0_ being determined by the ‘helical twisting power’ (HTP) of the chiral dopant and its concentration [*c*]: 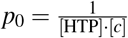^17^. While characteristic helical ordering and textures can become evident even with low concentrations of added dopant^4,18^, high dopant concentrations (between 25% and 35% w/w) are usually necessary to give rise to chiral phases with strongly reflected colours in the visible range^14,19–21^. These reflected colours, arising due to Bragg reflection, are a function of their helical arrangement and the viewing angle, analogous to the structural colours found in bird wings and in certain fruits^22^. Another notable aspect of CLCs is that, because the obtained colours are not solely due to birefringence, one can observe these striking colours without the aid of polarizers. This has the advantage of CLCs being more readily incorporable into an eventual device usable by lay technicians.

One of the oldest uses of CLCs in sensing is in quick-response thermometers, where changes in colour correspond to a change in temperature^23^. Several other sensors have incorporated CLCs in order to detect the presence of volatile organic compounds^14^, for pH^21,24^, for the detection of antibody-antigen binding events^25^, and into rubbers for strain sensing^26^. Less studied, however, are the effects of adsorption of different types of biomolecules to an CLC interface, and in particular the use of CLCs for sensing of surface active agents, especially amphiphilic biomolecules such as fatty acids and lipids, molecules which are key biological components of the (intra)cellular membrane, metabolic pathways, and which play a crucial role in various medically relevant conditions^2,27,28^ as well as possible contaminants in biodiesel production^15^. Using CLCs to sense these is a topic that has been touched on by H.G. Lee *et al*^21^ where they employed CLC droplets to detect synthetic surfactants, the addition of an amphiphile giving striking visible differences between a uniformly oriented structure and a multidomain texture. However, the use of CLCs for biological sensing is a field with much untapped potential for exploration.

In this paper, we show that micron-sized CLC droplets, selectively and sensitively, optically respond to amphiphiles in *in vitro* as well as *in vivo* settings, underlining their potential for rapid biological sensing. By exposing CLC droplets (*p*_0_~650 nm) to different species (surfactants, fatty acids, and lipids) and concentrations (order of *μ*M to mM) of amphiphiles, we observe clear optical differences with each species, providing a unique ‘optical signature’ that allows us to distinguish between molecules with the exact same carbon tail but differing headgroups. In order to make our sensing platform robust, easy-to-use, and easily transportable, we stably deposit the CLC droplets deposited on a glass surface at high densities in the dried state. These compact arrays of CLC droplets elicit the same optical responses when rehydrated with an amphiphile-containing sample, with the response time being as short as seconds for higher concentrations. Lastly, we demonstrate proof-of-principle *in vivo* testing by injecting the droplets in live zebrafish, showing a differential response for a healthy and inflamed gut environment.

## Results

### CLC droplets can sense diverse amphiphiles through distinct optical responses

We started with comparing the response of NLCs and CLCs, suspended in aqueous solutions, to surface active agents. We used pure 4-cyano-4’-pentylbiphenyl (5CB) and 35% CB15 in the eutectic nematic LC mixture RO-TN 407 to respectively produce NLC and CLC droplets; the main chemicals used in this work are shown in Figure S1. The size of our droplets was in the range 14.3–87.4 *μ*m (34.5±12.6 *μ*m; mean ± standard deviation). In order to stabilize the droplets and prevent their coalescence (owing to the considerable LC-water interfacial tension^16,29^), we used 87-89% hydrolyzed poly(vinyl alcohol) (PVA), a polymer typically used to stabilize LC shells and droplets while still preserving the anchoring conditions provided by pure water^30^. We therefore use PVA solution to suspend our droplets in future stages because it gives the same alignment as pure water. As can be see in Figure 1(a), NLC droplets adopted a homeotropic/normal configuration upon exposure to SDS, fatty acids (lauric acid, LA), and phospholipids (such as 1,2-dioleyoyl-*sn*-glycero-3-phosphocholine, DOPC): the typical planarly aligned texture, with surface defects imposed by the geometry (per the Poincaré-Hopf Theorem, requiring a minimum of two defects at the poles^31^), evolved into a typical ‘Maltese cross’ with a single, prominent point defect at the center (governed by the hairy ball theorem). This is a striking contrast but an expected outcome^10,32^. Using fluorescent lipids further confirmed that homeotropic switching was caused by lipid molecules adsorbing at and thus stabilizing the interface (Figure S2). However, irrespective of the amphiphile, as seen in Figure 1(a), the resulting droplet images between crossed polarizers were virtually indistinguishable.

**Figure 1.**
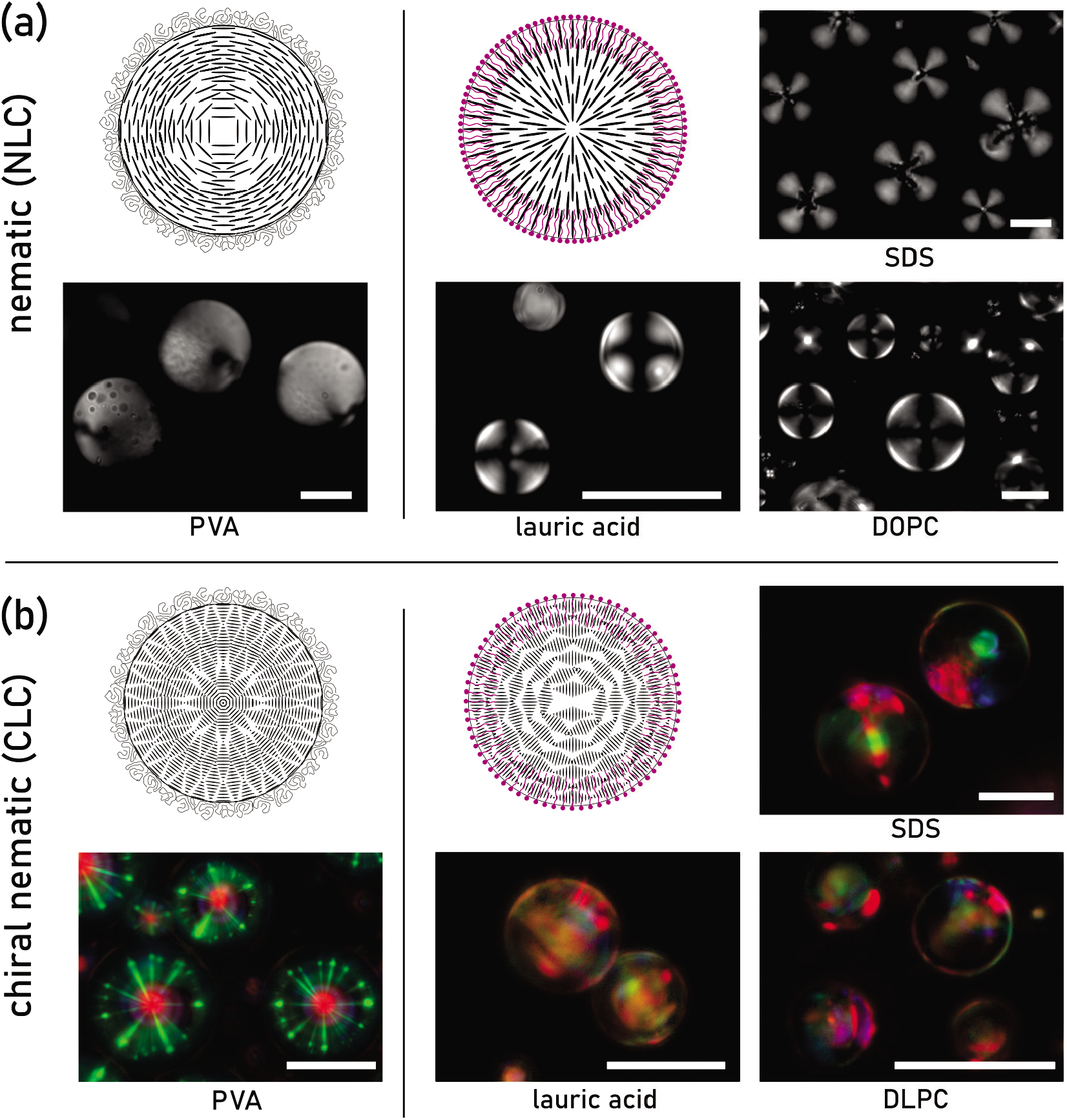
Sensing of amphiphiles using CLCs gives distinct reflected colours. Example amphiphile sensing using (a) a nematic liquid crystal and (b) a red-reflecting chiral nematic liquid crystal with corresponding micrographs. Samples were prepared in 0.2% w/w PVA solution and in solutions of a synthetic anionic surfactant (6.0 mM SDS), a long-chain fatty acid (5.0 mM LA), and phospholipids (40 *μ*M DOPC or 100 *μ*M DLPC). The NLC in (a) shows a difference from a planar/tangential structure in the presence of PVA (without amphiphiles) to a homeotropic/normal alignment, the hallmark of which is the large central Maltese cross. On the other hand, the additional helical ordering in the CLC in (b) shows a contrast from a starburst pattern with strong inter-droplet reflection to a polydomain, strongly scattering texture with oily streak defects across the surface in the presence of amphiphile. The multidomain texture is found to differ considerably based on the identity of the amphiphile added, while the micrographs in (a) show minimal difference, if any, based on the quality of the amphiphile added. Micrographs in (a) are obtained in transmission mode between crossed linear polarizers; micrographs in (b) were acquired in reflection mode with crossed linear polarizers. Scale bars 25 *μ*m.

The scenario changed significantly when we used CLCs to sense the same amphiphiles. In the absence of a surfactant, the PVA-stabilized droplets exhibited a starbust pattern, composed of a single red-reflecting point corresponding to Bragg reflection from the equilibrium cholesteric texture surrounded by green rays resulting from a blue-shifted Bragg reflection between droplets (Figure 1(b)). In samples with very few droplets, the intensity and number of inter-droplet reflected rays is greatly reduced, confirming that the green ‘rays’ are a direct consequence of inter-droplet reflection (Figure S4). The adsorption of amphiphile at the interface, however, transitions the alignment from the uniform texture with the helix oriented normal to the droplet surface to a scattering, polydomain texture; this is imposed by the alignment switch induced by the adsorption of the amphiphile combined with the continued helical modulation, which creates a situation akin to a rope wrapped around a sphere.

The CLC droplets were remarkably sensitive to amphiphiles, responding to samples with concentrations on the order of *μ*M of SDS, LA, and DLPC (Figure S3); similar sensitivity was observed in NLCs. This is in striking contrast to previous works which sensed amphiphiles at much greater concentrations^11,21^. More importantly, in case of each amphiphile, a distinct reflected colour profile was visible, especially compared to the uniform Maltese crosses observed in case of NLC sensors: the starburst pattern transitioned into a much less uniform, more scattering texture with defects and sometimes multiple bright reflecting spots. The green in the reflection often became dominated by red or other colours. Observing these qualitative differences was an encouraging sign to further investigate the potential of CLCs in sensing amphiphiles.

### CLC droplets can be dried and rehydrated to quantitatively sense amphiphiles

Once it was clear that CLC droplets could sense amphiphiles, giving distinct reflected colour profiles, we proceeded with quantifying the sensing process. At the same time, we also wanted to make it easier to visualize and render the assay easily transportable. Instead of having randomly distributed CLC droplets in a three-dimensional space, a dense quasi-2D array would satisfy both these requirements. To achieve this, we attempted to air-dry PVA-stabilized droplets on a glass slide. For the drying process, we chose a partial degree of hydrolysis (87-89%) because intermediate grades of hydrolysis are more amphiphilic than lower or higher grades and the amphiphilicity is necessary to create the shell of PVA that will protect the droplets during the drying process^33^. We observed that not only the droplets were completely preserved through the drying process, but they formed highly compact arrays and gave the expected signal when rehydrated, as shown in Figure 2. Attempting to dry droplets without PVA was unsuccessful in preserving droplets due to the collapse of droplets onto the glass and wetting. We then exposed the dried droplet arrays to aqueous solutions containing a target amphiphile, which resulted in a species-specific reflected colour response, as presented in Figure 2. We performed the drying-rehydration procedure for three different amphiphiles: SDS (0.6 mM; Figure 2(e)), LA (1.0 mM; Figure 2(f)), and lipid A (1.0 mM; Figure 2(g)), a glycophospholipid component of a bacterial endotoxin with six lipid tails. Rehydration with PVA solution (0.2% w/w; Figure 2(d)) acted as a negative control. We quantified the optical response by obtaining the average intensities of the primary colour channels (red, green, and blue) of individual droplets. In order to eliminate the influence of photography parameters such as background illumination, exposure time, and light intensity, we analyzed the intensity ratios (R/G, G/B, R/B) to study the differences between the responses generated from different amphiphiles (see Methods for details). This straightforward analysis gave us a unique ‘optical signature’ for each amphiphile, as shown in Figure 2(d-g). For each of the four environments, at least one colour ratio showed a statistically significant difference with all the other three amphiphiles. To begin, owing to the low values of the R/G (0.79) and R/B (0.97) ratios, the sample in PVA (Figure 2(d)) showed significant differences from all the other samples for these two ratios in particular. In the case of SDS, the R/G (1.17) and R/B (1.42) ratios showed no significant differences with LA (1.11 and 1.49), but the differences in the G/B ratio (1.21 versus 1.32) were statistically significant (Figure 2(e)). The G/B ratio is a good signature for LA (1.32) because this combination was significantly different compared to SDS and PVA groups, whereas the R/G and R/B values statistically diverged when compared to PVA and Lipid A (Figure 2(f)). For Lipid A, the R/G (1.78) and R/B (2.22) ratios were significantly higher than all the other groups, thus undoubtedly serving as its unique signature (Figure 2(g)). This information is summarized in Figure 2(c). Our analysis demonstrates that we can effectively use colour response to quantitatively detect the difference between homologous amphiphiles with different headgroups.

**Figure 2.**
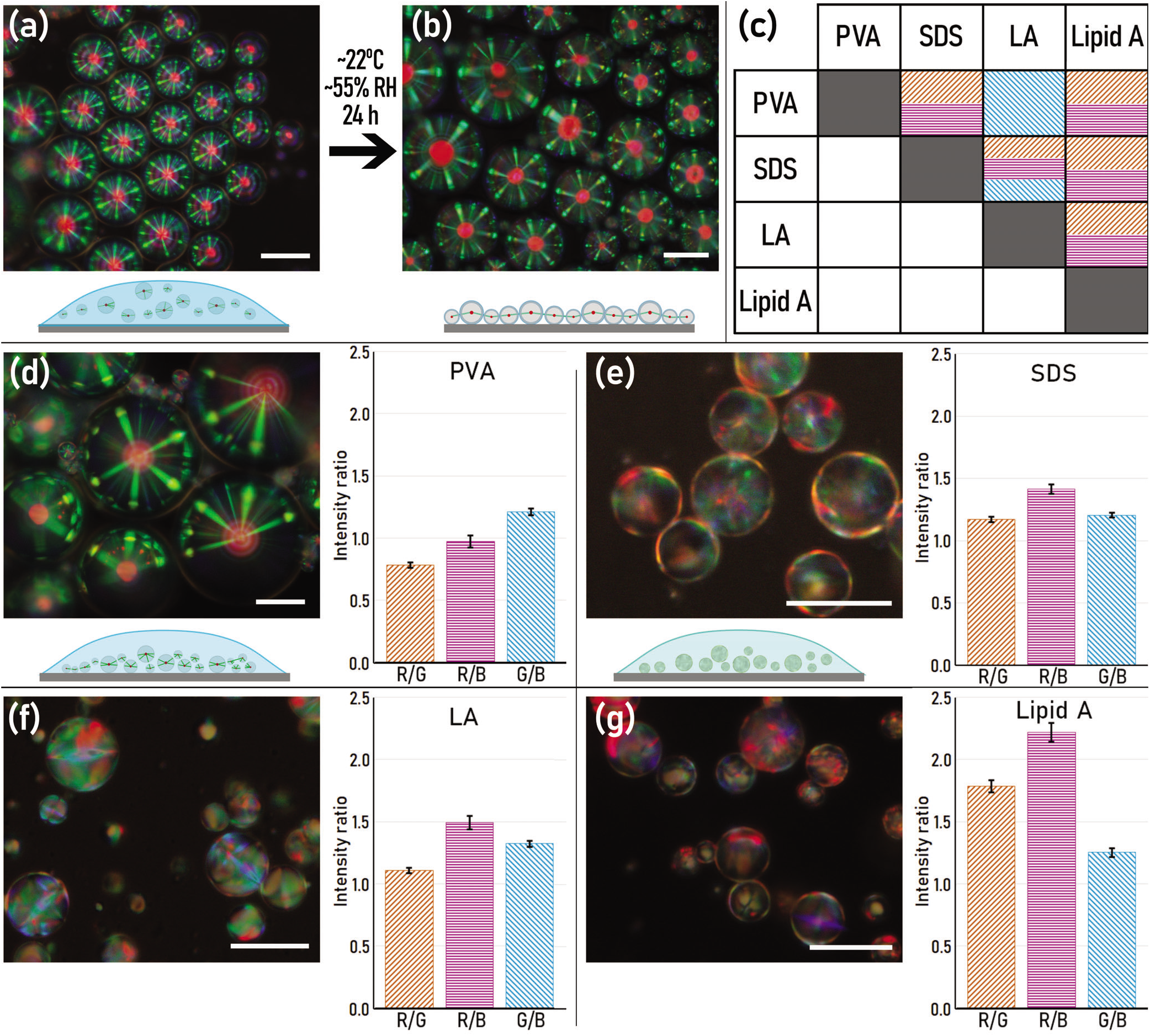
CLC droplets can be stably dried in a responsive state and readily react upon introducing amphiphiles. (a) Droplets of the CLC were produced in a 0.2% w/w PVA solution in Tris buffer, stabilizing them, and then deposited on a glass slide. Strong inter-droplet reflections in the form of a green starburst pattern due to the normal helical alignment and tangential LC alignment are clearly observed. (b) By drying the deposited solution under ambient conditions (22°C) overnight, the CLC microdroplets, coated with PVA, are stabilized and maintain their shape in the dried film, while still preserving the characteristic starburst reflection associated with tangential LC/normal helical alignment. (c) An analysis matrix showing that the ratios of the primary colour intensities (R/G in brown, G/B in magenta, and R/B in cyan) can be used to distinguish between different amphiphiles. Each coloured box indicates a comparison for which *p*≤0.01. (d) Rehydrating the slide with PVA buffer solution preserves the alignment of the LC droplets (e-g) and rehydrating the film with an amphiphile buffer ((e) 0.6 mM SDS; (f) 1.0 mM LA; (g) 1.0 mM Lipid A) solution will reorient the CLC, with the intensity ratios of the three channels dependent on the amphiphile used. The summaries of the relationships of these ratios, as shown in (c), show these can be used to assign a unique signature for each situation. Scale bars 25 *μ*m. Error bars represent standard error of the mean.

We further analyzed whether the optical signature of an amphiphile varied across the concentration range (Figure S5). First of all, switches in colour were apparent even at extremely low concentrations of amphiphiles, as shown in Figure 3(a) where 1 *μ*M SDS was sufficient to switch the alignment of the CLC, producing a clear response. In case of SDS, we were able to distinguish between three concentrations: the 6 mM and 12 mM groups showed a statistical difference in B/R and G/R, respectively. In case of LA, no statistically significant differences found over the small concentration range (1-5 mM) that was tested. These results demonstrate further potential in sensing the amphiphile concentration based on the reflected colour. However, what we found to be even more effective was the response time, i.e., the time required after rehydrating a sample to fully undergo switch of alignment from the tangential/planar case to the normal/homeotropic condition (see Methods for details). We noticed clear differences in the switching times upon amphiphile exposure (SI Videos 1-3 and Figure 3(b)): upon increasing the concentration of amphiphile, the response time to switch the droplets greatly decreased from over 2 min at 0.6 mM SDS to under 10 s at 12 mM.

**Figure 3.**
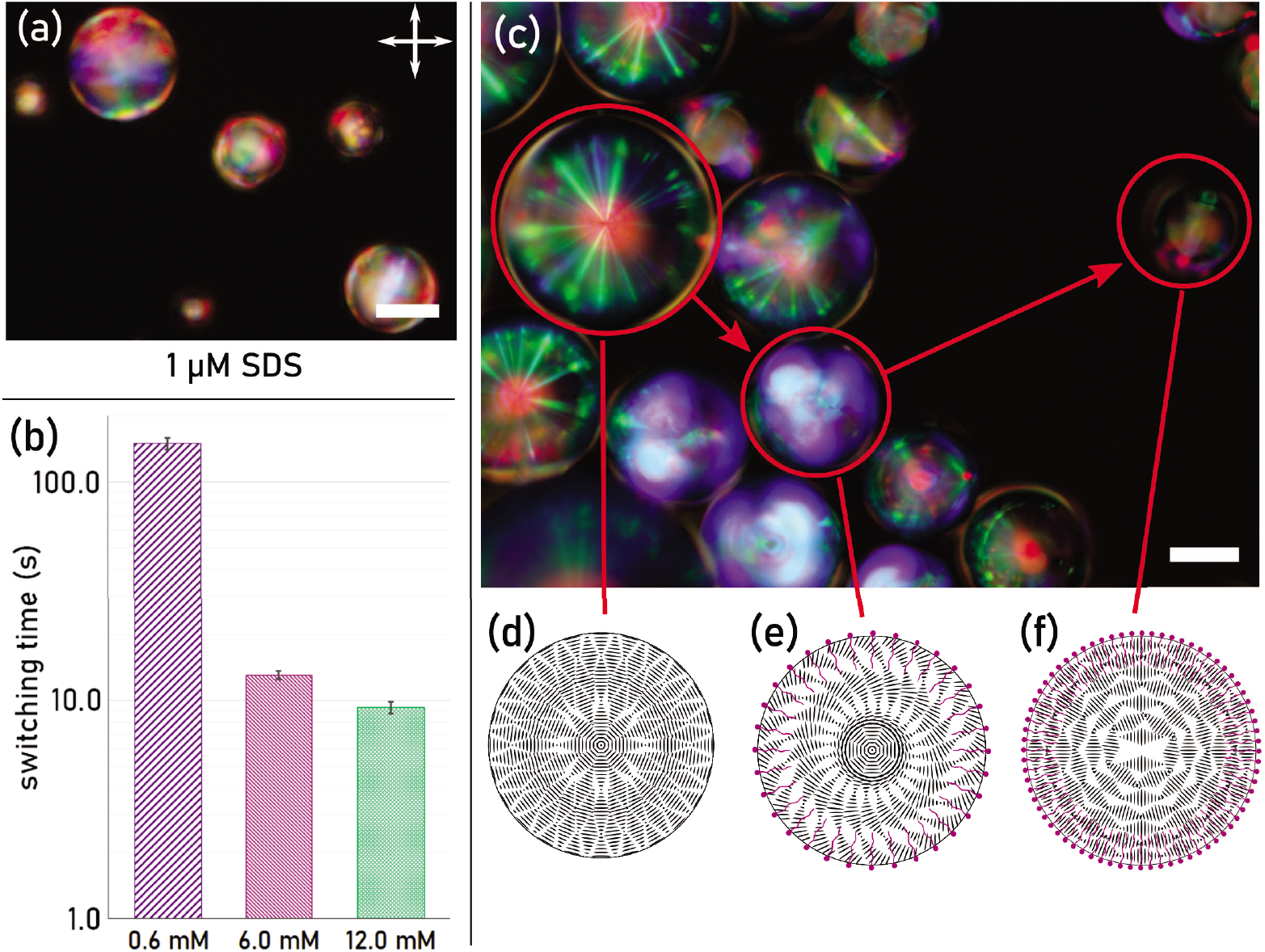
CLC droplets can detect amphiphiles at low concentrations, have fast response times, and exhibit complex switching dynamics. (a) A sample prepared from 1 *μ*m SDS in Tris buffer, showing switching of the droplets at very low concentrations of amphiphile. Scale bars are 25 *μ*m. (b) By exposing a dried array of droplets to different concentrations of an amphiphile (here, SDS), we observe marked differences in the switching times, ranging from 9.3±0.5 s for 12.0 mM SDS solution to 150±9.5 s for 0.6 mM SDS solution. Analysis based on the switching times of at least 20 randomly selected droplets in each batch. Error bars indicate standard error of the mean. (c) A sample of CLC droplets exposed to 5.0 mM lauric acid in Tris buffer showing the different stages of the switching process, ranging from (d) fully tangentially/planarly aligned, where a single red reflection and a starburst pattern from inter-droplet Bragg reflection is evident; (e) a transient metastable texture, where the helix axis is tilted off-vertical, producing a blue-shifted Bragg reflection; and (f) fully normally/homeotropically aligned, where we instead see a polydomain reflection without a single, prominent central reflection. Viewed between crossed linear polarizers.

Figure 3 (c-f) shows the switching process in a sample exposed to lauric acid, showing several droplets at different stages of the transition between alignments, corresponding to SI Video 4. As the droplets begin to switch from tangential/planar alignment to normal/homeotropic alignment, several features become noticeable. The equilibrium tangential texture, with a single red reflection accompanied by a green ‘starburst’ pattern, is due to Bragg reflection from the CLC helix from different angles of incidence. Top-down illumination, corresponding to an illumination at an angle of 0°, produces a single red central reflection spot, which we expect due to the CLC being prepared to reflect red, while inter-droplet Bragg reflections created from illumination at greater angles will produce the green starburst pattern. The use of different CLC samples with different equilibrium pitch values would shift the colours reflected, but the principle will remain the same, with the inter-droplet reflections being blue-shifted compared to the central reflection.

The dynamics of switching of the LC droplets are multifaceted, especially in comparison to thin films, with the ultimate alignment a complex interplay between surface and bulk effects. During the switching process, one of the first things to change is that the droplets will briefly appear blue (Figure 3(c)). This is an expected response due to the LC realigning: since the helical orientation begins to tilt from the outside inwards as the alignment switches, the wavelength of light that satisfies the Bragg condition becomes blue-shifted^19,20^ as the angle of incidence becomes off-vertical. As dictated by the Poincaré-Hopf theorem, the helical modulation of the CLC in a normal/homeotropic anchoring condition cannot smoothly lie along the surface of the droplet, but requires at least two defects in the orientation. The nature of the switching process and the degeneracy of the anchoring condition means that a large number of defects, forming a metastable state, can develop during the switching process. These defects and inhomogeneities in the LC structure thus lead to the complex, multidomain texture observed in the droplet reflections.

The energy landscape of a given material with a director field represented by the unit vector 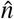, a director field representing the preferred LC orientation, is a combination of the elastic deformation energy, described using the Frank–Oseen free energy density^32,34^ 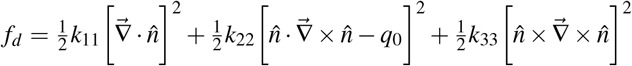, where *k*_11_, *k*_22_, and *k*_33_ are elastic deformation constants corresponding to splay, twist, and bend of 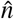, respectively, and *q*_0_ is a term referring to the equilibrium pitch of a CLC; and the surface energy associated with the interfacial tension. Generally, for thermotropic liquid crystals (such as 5CB), *k*_33_ > *k*_11_ > *k*_22_^35^. This means that the normal/homeotropic anchoring case is favoured in a droplet; however, the surface energy associated with normal/homeotropic anchoring (*γ*_⊥_) is higher than for tangential/planar anchoring (*γ*_∥_)^16^. The total energy is thus described by a combination of both the surface and the volume terms. A full analytical treatment is beyond the scope of this paper; however, the effects of this are that, since *γ*_∥_ < *γ*_⊥_, but *k*_33_ > *k*_11_, there is a threshold of surfactant or amphiphile adsorption necessary to reduce *γ* so that the energy of the splay configuration is more favourable. Often this interfacial tension decrease is sufficient at very low concentrations of surfactant (on the order of *μ*M^16^) to induce switching, as shown in Figure 3(e). We notice the effects of this in terms of switching time, with smaller droplets switching somewhat more quickly. This is most likely due to a lower overall energy necessary for switching, but we also observed that not all droplets of the same size switched simultaneously, indicating that spatial positioning plays a role. We further confirmed that droplet size did not ultimately play a significant role on the observed optical response. As Figure S6 shows, the colour ratios (here, in case of lauric acid) for different droplets grouped on the basis of similar diameters gave very similar values, indicating that the optical response was not affected by the size of the CLC droplet, at least within the size range of our droplets.

### CLC droplets can be used for *in vivo* sensing with zebrafish

With the above encouraging results, we further probed whether these droplets could be actually used as *in vivo* biosensors. To test this idea, we decided to use zebrafish as our model system. The zebrafish *(Danio rerio)* is often used as a model vertebrate organism for the study of many processes owing to its well-sequenced genome, fast development, and well-understood developmental behaviours. We chose to experiment with 5 day-old larva (5 days post-fertilization (5 dpf)), as their entire intestinal tract is fully open and functional at that point. Additionally, the zebrafish being optically transparent facilitates the visualization of the LC droplets both in transmission and reflection mode.

While the LC materials we use are themselves cytotoxic^36^, we figured that coating our LC droplet with PVA would make them fairly biocompatible. In Figure 4, we illustrate how CLC droplets can be used to sense the presence of biologically relevant amphiphiles in the gut of a zebrafish. We embedded the fish in an agar gel and gavaged them with 4 nl of the CLC droplet suspension (Figure 4(a)). Remarkably, the droplets lodged in the gut (Figure 4(b-d)) showed a complete lack of the central red reflective spot (indicative of the default planar alignment), but instead exhibited a diffuse reflection, associated to a switch because of surface active agents such as amphiphiles (Figure 2). The green signal we observe is similar to the ‘chicken skin’ pattern and texture described by H.G. Lee et al^21^ that was present when amphiphiles were adsorbed to the interface. We saw that this reflection was much less complex in terms of texture compared to the *in vitro* sensing experiments, as the multidomain scattering texture we would typically see is not present, though this may be an effect of the difficulty of imaging inside of the zebrafish. Furthermore, due to the transparency of the zebrafish larvae, the reflected signal was observable even without the use of polarizers (Figure 4(c)). However, the signal does become cleaner and more distinctive with polarizers (Figure 4(d)), as the polarizers cut out any reflection from non-birefringent materials. Thus, we can conclude that amphiphilic molecules present in the gut, likely the metabolic byproducts of the gut microbiota^37,38^, are responsible for switching the droplet alignment. The aqueous medium in which the fish were swimming did not switch the droplet alignment. A much better control was observed in the form of the LC droplets lodged within the oral cavity, clearly showing a single central red reflection (Figure 4(f)), which is characteristic of the reflection from droplets without adsorbed amphiphiles (Figure 4(e)). This further confirms that the droplet switch observed within the gut is indeed sensing the amphiphilic compounds being produced within, most likely medium-to long-chained fatty acids, rather than lipids, as the concentrations of lipids necessary to induce a response in PVA-coated droplets *in vitro* was found to be typically high. We also observe that the cartilage and bones of the zebrafish are, in fact, birefringent, much like we would expect from any aligned long-chained polymer or crystalline structure^20^, but this birefringence is of much lower order than the LC^39–41^ (Figure S7). Many of the melanocytes also appeared birefringent, but their signals were readily distinguishable from the signals from the CLC droplets (Figure S7).

**Figure 4.**
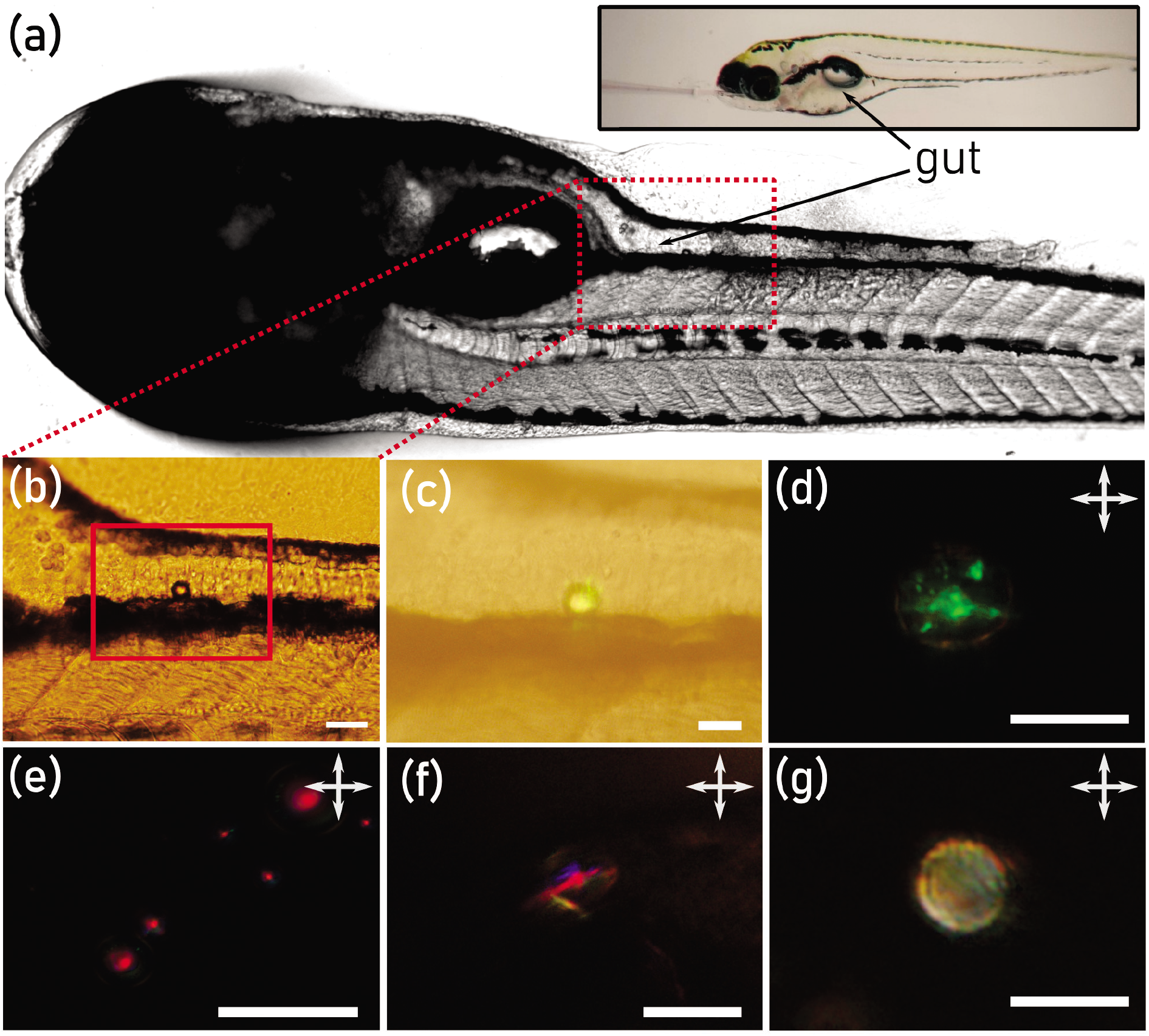
CLC droplets can detect amphiphiles within the zebrafish gut. A zebrafish at 5 days post-fertilization, gavaged with a dispersion of the CLC mixture suspended in a 0.2% w/w low-weight PVA in Tris buffer solution. (a) The zebrafish, with its gut clearly visible, viewed with a stereomicroscope. Inset: a demonstration of the gavaging process, with the glass capillary inserted into the oral cavity. (b) An inset showing a CLC droplet that has been lodged in the gut. (c) By using reflection mode microscopy without polarizers, we can see the droplet reflecting a green colour. (d) Close-up in reflection mode microscopy between crossed polarizers clearly shows that the droplet no longer has the characteristic starburst texture associated with a planarly-aligned CLC droplet and has a diffuse, scattering texture. This indicates the presence of an amphiphile, with the polarizers cutting out reflection from non-birefringent surfaces. (e) An *in vitro* control showing the characteristic red reflected central spot present when no amphiphiles are adsorbed to the LC interface. (f) The same characteristic central red spot seen in a droplet residing in the oral cavity of the fish instead of in the gut. (g) The use of soyasaponin, a chemical known to induce intestinal inflammation, shows a strikingly different colour pattern compared to (d), suggesting the presence of something different. Scale bars in (b), (e), and (f) 50 *μ*m; (c), (d), and (g) 25 *μ*m. (d)-(g) viewed between crossed linear polarizers.

Finally, we wondered whether we could see a difference between a healthy and inflamed gut by recording the optical response of LCs. For that, we grew the zebrafish larvae in a solution containing 0.5 mg/ml soyasaponin, an inflammatory agent^42^. Upon gavaging these treated fish with CLCs, we again saw a lack of the central red reflection, but, most remarkably, the texture of the CLC droplet appears markedly different from that observed in case of healthy fish (Figure 4(g)). These proof-of-principle experiments clearly show CLC droplets show great potential for *in vivo* sensing.

## Discussion and Conclusion

We have shown that cholesteric liquid crystals can be efficiently used to detect biorelevent amphiphiles such as fatty acids and phospholipids. We were able to detect analytes at *μ*M concentrations: this corresponds to ng/ml concentrations in case of lipid A and is similar to the limits of detection observed previously^43^. We systematically quantified the optical response of different amphiphiles and showed that a simple parameter, *i.e*, ratio of the primary colour channels, is sufficient to assign a unique signature to different amphiphiles. With the detection of amphiphilic analytes in portable on-chip settings as well as in highly complex biological environments such as the intestinal tract of zebrafish larvae, our easy-to-implement CLC sensors show high potential in biosensing.

We see several ways of further improving our sensors. A priority would be improving the detection limit of the dried droplet arrays. In the current state, the concentrations of amphiphiles necessary to induce an optical response in dried sensor arrays were often quite high, on the order of mM. This is not the case for bulk samples, where *μ*M concentrations were enough to induce alignment changes. A very likely cause of this is the presence of PVA at the droplet interface. A small amount of PVA was necessary to prevent the droplets from wetting the glass slides onto which they are deposited and also to prevent coalescence with each other. The protective coating formed by the PVA polymers around the LC droplets impedes the adsorption of amphiphiles. We used a relatively high PVA concentration of 0.2% w/w, corresponding to ~ 100 *μ*M, in order to stabilize the droplets during the drying process. Additionally, PVA molecules are considerably larger compared to the size of the amphiphiles detected (~60× larger), further shielding the CLC droplets from amphiphile exposure. Lower concentrations of PVA were tested, but often did not keep enough droplets stable during the drying process for eventual sensing.Other non-surfactant polymer stabilizers may have a better outcome in giving us the best of both worlds (high sensitivity of amphiphile detection while sufficiently stabilizing them for the drying process), a topic for future investigation. The reduction of sensitivity when using PVA, on the other hand, could also be used as a useful feature, setting the sensitivity of the LC to a specific level to function as a threshold sensor, detecting the target analytes only above a specific concentration.

Despite the cytotoxicity of many LC mixtures^36^, the CLC droplets did not appear to have an adverse effect on the health of the zebrafish; the adverse effects came instead from the embedding in agarose for the long duration (few hours) of the experiments. Nevertheless, for future work, it is conceivable that biological analogues that form LC phases, such as cellulose nanocrystal dispersions^44^, or more biologically friendly CLC generating materials such as cholesteryl ester mixtures^45^, could be used as a more biocompatible sensing material. The capability to sense amphiphiles *in vivo* within the zebrafish additionally shows great promise in being able to rapidly detect and sense the presence of biologically relevant markers, such as amphiphilic short-chained fatty acids, which typically require complex set-ups or sophisticated equipment to detect^38^. Other applications for *in vivo* sensing in fish include detecting how well feed is uptaken and metabolized, enabling for further optimization of animal husbandry^37^. We could envision incorporating CLC droplets into the feed, either for imaging within the animal or for post-analysis after passing through the digestive tract; such an ingestion-and-retrieval strategy of microsensors to detect and profile the interacting bioamphiphiles may give the opportunity to gain information about the health of animals much more rapidly and potentially capable of used by lay technicians without the necessity of specialized equipment.

In conclusion, we have demonstrated the potential of CLC droplets for biological sensing, both *in vitro* and *in vivo.* There is a wide scope in terms of what can be detected, automation in signal analysis, and into what novel platforms we can incorporate CLCs, such as simple fatty acid biosensors compared to those based on genetic modifications^46^ and enzymatic reactions^47^. Our future work will look in several directions to test the limits of CLCs for sensing, both in terms of qualitative and quantitative analysis; the use of CLCs when incorporated into specific immunoassay sensors^28^; to produce the droplets in a high-throughput, controlled (e.g., to obtain monodisperse samples) fashion using microfluidic systems^48^ and further store them efficiently to improve detection^49,50^; and in examining cooperative effects in terms of specificity.

## Methods

### Materials

The nonchiral nematic liquid crystal 4-cyano-4’-pentylbiphenyl (5CB, 95%+, Figure S1(a)) was obtained from Tokyo Chemical Industries. The chiral nematic liquid crystal mixture RO-TN 407 (F. Hoffmann-La Roche) mixed with 35% w/w CB15 ((*S*)-4-cyano-4’-(2-methylbutyl)biphenyl, Synthon GmbH, Figure S1(b)), producing a red-reflecting LC mixture (*λ* ~ 650 nm), was graciously provided by the University of Luxembourg. Poly(vinyl alcohol) (PVA) in two weights (high weight: *M_w_* = 30-70 kDa, 87-89% hydrolyzed; low weight: *M_w_* = 13-23 kDa, 87-89% hydrolyzed), sodium dodecyl sulfate (SDS, 99%+, Figure S1(d)), and dimethyloctadecyl[3-(trimethoxysilyl)propyl]ammonium chloride (DMOAP, 42% solution in methanol) were obtained from Merck–Sigma Aldrich. Trizma base (tris(hydroxymethyl)aminomethane), and Trizma HCl (tris(hydroxymethyl)aminomethane hydrochloride) were sourced from Sigma Aldrich and combined in ultrapure water to obtain a buffer with pH 7.4 (Tris 7.4). Lauric acid (dodecanoic acid, LA, 99%, Figure S1(c)) was obtained from Acros Organics. The phospholipids DOPC (1,2-dioleyoyl-*sn*-glycero-3-phosphocholine, 25 mgoml^-1^ in chloroform), Liss-Rhod DOPE (1,2-dioleyoyl-*sn*-glycero-3-phosphoethanolamine-N-(lissamine rhodamine B sulfonyl) (ammonium salt), Figure S1(f), 1 mgoml^-1^ in chloroform), and DLPC (1,2-dilauroyl-*sn*-glycero-3-phosphocholine, Figure S1(e), 25 mg·ml^-1^ in chloroform) were sourced from Avanti Polar Lipids. Kdo2-Lipid A (di[3-deoxy-*D*-manno-octulosonyl]-lipid A (ammonium salt), Figure S1(h), lyophilized powder, 90%+) was purchased from Merck-Sigma Aldrich. Soyasaponin extract (95%, Figure S1(g)) was obtained from Organic Technologies (Ohio, USA). Ultrapure deionized water (resistivity 18.2 MΩocm) was purified with a MilliQ system.

### Solutions

We prepared stock solutions of 0.2% w/w PVA (of both weights), 6.0 mM and 12.0 mM SDS, and 5.0 mM LA in Tris 7.4 buffer and allowed them to mix at room temperature until the powders were fully dissolved, as verified by the presence of an optically transparent solution with no lumps or aggregates.

For phospholipids, appropriate volumes of lipids in chloroform were pipetted into a clean glass vial. Chloroform was then evaporated under vacuum until the films were dried, after which the lipid vials were hydrated with the buffer solution at room temperature for at least 4 hours. Lipid dispersions were either then sonicated using a pulsed tip probe sonicator (10% power, 0.1 s on/0.9 s off) for 5 min or extruded twice through a 0.22 *μ*m PTFE syringe filter into an Eppendorf tube to obtain a clear, optically transparent dispersion. In the case of Kdo2-lipid A, we directly massed the appropriate quantity of powder into an Eppendorf tube before adding an appropriate quantity of buffer solution, agitating, and filtering to obtain a clear dispersion. All solutions were used within a week of preparation.

### Protocols

#### Droplet Visualization, Drying, and Rehydration

We visualized the LC droplets both in bulk solutions and when dried on glass slides. For bulk LC droplet samples, we pipetted 250-1000 *μ*l bath solution along with 2.5-5.0 *μ*l LC solution in a clean 1.5 ml Eppendorf tube, in order to obtain a dilute dispersion of droplets. This dispersion was alternately vortex-mixed at a high power for 30 s and manually shaken to obtain a cloudy dispersion. We then immediately pipetted 10-20 *μ*l of dispersion onto a clean glass slide or into a polydimethylsiloxane (PDMS) well covalently bonded to a glass coverslip through plasma bonding.

To prepare microscopy slides with dried arrays of LC droplets, fresh glass slides were either simply rinsed with MilliQ water and air dried with a compressed air pistol or silanized by plasma cleaning using a Harrick PDC-32G plasma cleaner/sterilizer for 60 s followed by immediate immersion in a dilute solution of DMOAP (1% v/v DMOAP in ultrapure water) for 30 min, with gentle shaking throughout to ensure all glass surfaces were adequately exposed to the surfactant. The treated slides were then washed at least three times with deionized water before baking for 4 h to overnight at 115°C in an oven. The success of the surface treatment was verified by a simple contact angle observation with ultrapure water to check for wetting.

Two to three droplets (10 *μ*l each), of the prepared LC droplet dispersion were deposited on each glass slide (hydrophobized or untreated) with a minimum separation of 10 mm between each droplet. The slides were allowed to dry overnight under ambient conditions (22°C, 55% RH). A 10 *μ*l droplet of analyte solution was then gently pipetted on top of the dried droplet array and continuously imaged with manual focus adjustments during video acquisition.

#### Image Analysis

The colour images of droplets in varied microenvironments were analyzed using MATLAB R2019b to estimate both the intensity values and the mutual ratios between RGB channels. The values of the three channels were defined as the average intensities of the respective channels for each droplet. One-way analysis of variance (ANOVA, *α =* 0.01) was used to confirm if there was any significant difference for the R/G, R/B, or G/B ratios between droplets treated by different amphiphiles. Error bars represent the standard error of the mean (n > 50 droplets for every sample). Response time was analyzed by selecting a minimum of 20 random droplets in a video and recording the timestamps at which switching began (typically indicated by a blue ‘flash’) to the time when a steady state was reached, the progression of which is illustrated in Figure 3(a).

#### In Vivo Experiments

Wildtype (AB) zebrafish were grown both in E2 buffer (5 mM NaCl, 0.17 mM KCl, 0.33 mM CaCl_2_, and 0.33 mM MgSO_4_) or from days 2-5 post-fertilization (dpf) in a E2 buffer solution containing 0.5 mg·ml^-1^ soyasaponin extract to induce intestinal inflammation. Zebrafish grown in soyasaponin were then washed with E2 buffer to remove excess saponin. Once they reached age 5 dpf, they were anaesthetized in 3-amino benzoic acid ethyl ester (Tricaine/ethyl 3-aminobenzoate; Sigma Aldrich; 168 *μ*g·ml^−1^ in Tris pH 7) and embedded in 1% low melting point agarose (UltraPure Agarose, ThermoFisher Scientific) in E2 medium containing a small amount of anaesthetic (168 *μ*g·ml^−1^ Tricaine). Zebrafish were orally gavaged^51^ with 4 nl of the CLC mixture using glass capillaries (1.0 OD × 0.78 ID × 100 L mm, Harvard apparatus) by means of a micromanipulator and Eppendorf FemtoJet set-up. Zebrafish were imaged using a Leica DM6 upright microscope and an Olympus polarizing optical microscope equipped with a DT73 colour camera.

## Acknowledgements

We thank Prof. Jan P.F. Lagerwall, Dr. Hakam Agha, Dr. Catherine G. Reyes, and Dr. Shameek Vats from the University of Luxembourg for graciously supplying the cholesteric LC samples used in this study and for useful discussions; the students of the 2021 Advanced Soft Matter Practical Class at WUR (Maarten Dols, Axel Eijffius, Liza Leijten, Marlene Vollmer, and Niels Wensink) for performing some of the preliminary studies associated with this work; Martijn van Galen, Rob de Haas, and Nicolò Alvisi for technical support; and Prof. Oleg Lavrentovich and Dr. Piotr Popov for stimulating discussions. S.D. acknowledges financial support by the Innovation Program Microbiology grant (IPM-3) and by a ENW-KLEIN grant (OCENW.KLEIN.465) from the Dutch Research Council (NWO).

## Author contributions statement

L.W.H, S.B., and S.D. conceived the experiments; L.W.H., S.B., F.D., and S.D. performed the experiments; L.W.H, C.C., F.D., S.B., and S.D. analyzed the results; and L.W.H., C.C., and S.D. wrote the manuscript with input from the rest of the authors.

