## Supplementary Information for "Designing Biological Micro-Sensors with Chiral Nematic Liquid Crystal Droplets"

### Chemical Structures

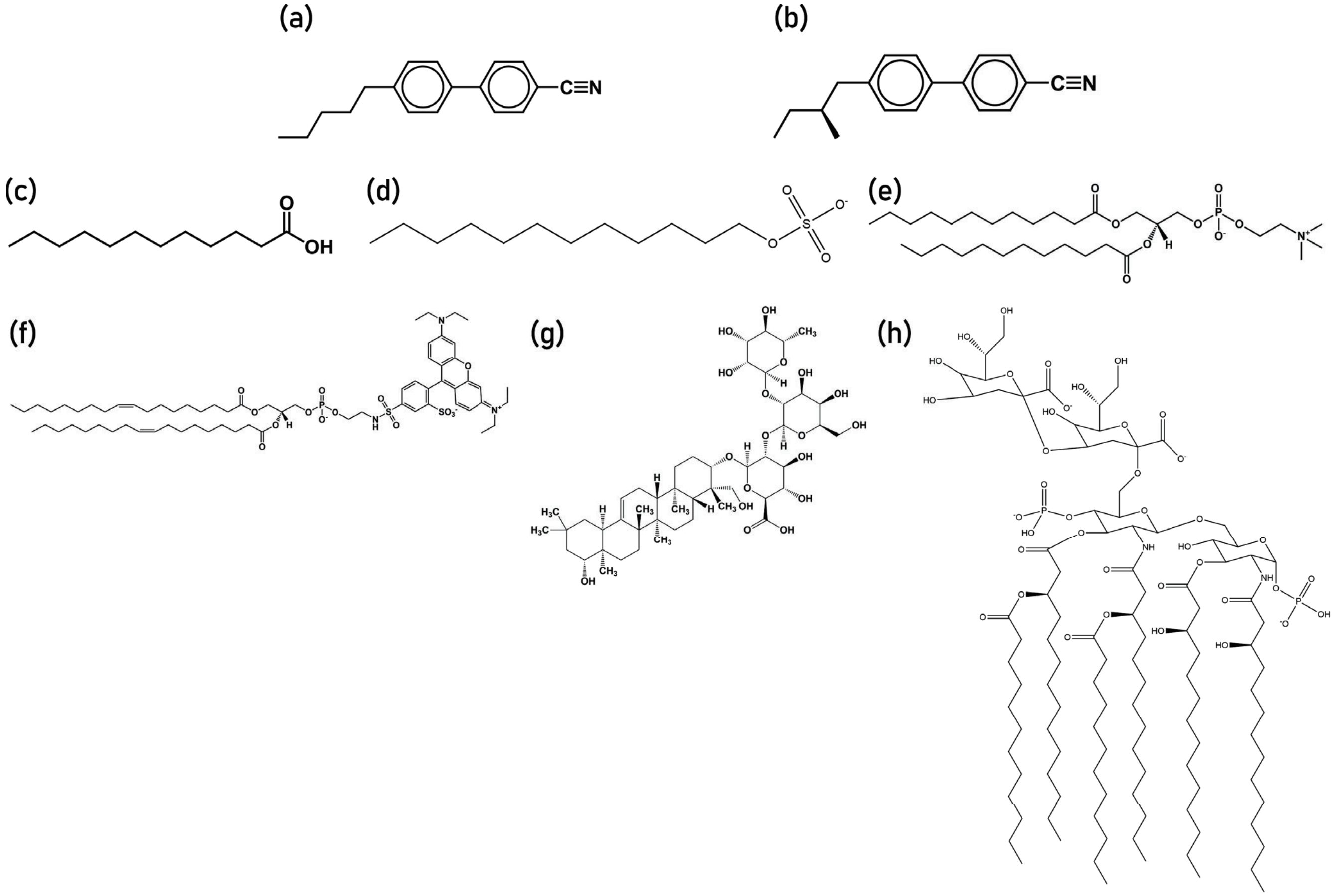


**Figure S1.** Structures of the main chemical compounds (liquid crystals, amphiphiles, and proteins) used in this work. (a) 4-cyano-4′-pentylbiphenyl (5CB); (b) (*S*)-4-cyano-4′-(2-methylbutyl)biphenyl (CB15); (c) dodecanoic/lauric acid (LA); (d) sodium dodecyl sulfate (SDS); (e) 1,2-dilauroyl-*sn*-glycero-3-phosphocholine (DLPC); (f) 1,2-dioleyoyl-*sn*-glycero-3-phosphoethanolamine-N-(lissamine rhodamine B sulfonyl) (ammonium salt) (Liss-Rhod DOPE); (g) soyasaponin; and (h) Kdo2-Lipid A (di[3-deoxy-*D*-manno-octulosonyl]-lipid A (ammonium salt)).

### Microscopy and Supplemental Data

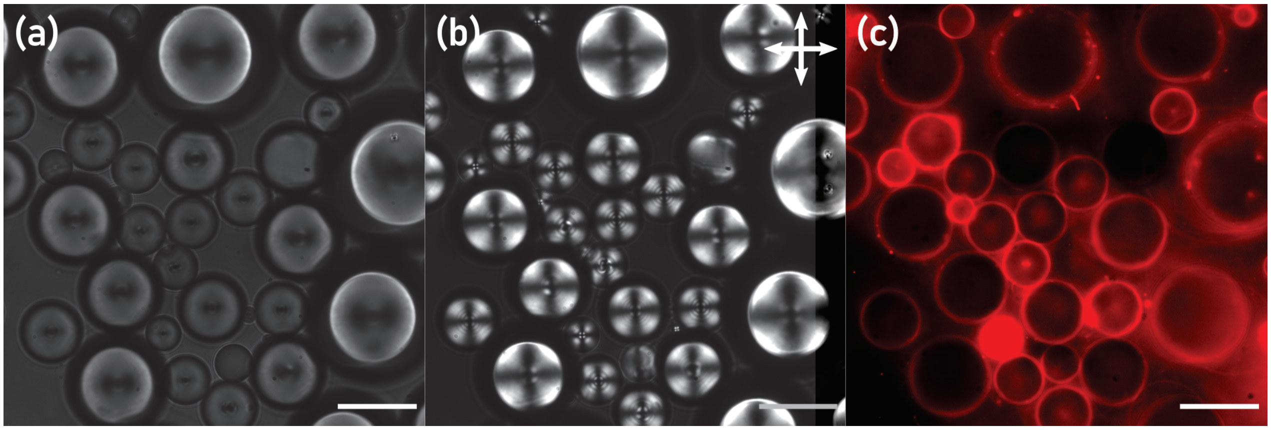


**Figure S2.** Droplets of 5CB prepared in a solution of 4 $\mu$M DOPC with 1:500 added Liss-Rhod DOPE in water, viewed (a) in transmission mode without polarizers; (b) between crossed polarizers, showing the characteristic Maltese cross textures of homeotropic-/normal-aligned NLC droplets; and (c) with epifluorescence, showing fluorescence signals corresponding to lipids adsorbed at the interface of the droplet. The adsorbed lipid is a proximal cause of the change of alignment. Scale bars 25 $\mu$m.

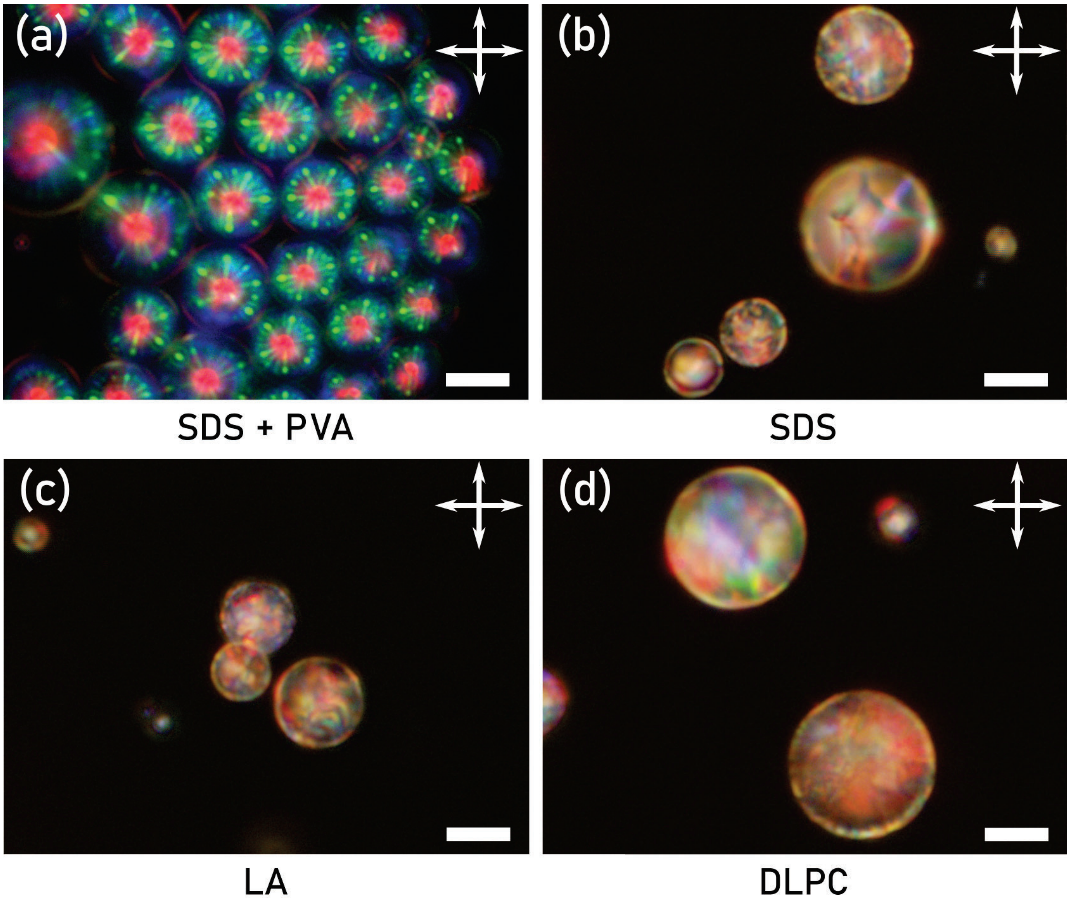


**Figure S3.** POM micrographs of a CLC sample (a) prepared in 0.2% PVA, dried, and rehydrated with 0.06 mM SDS solution, showing no difference from a sample containing just PVA; (b) prepared in 1 $\mu$M SDS solution without PVA; (c) in 10 $\mu$M LA solution; and (d) in 10 $\mu$M DLPC solution. The presence of PVA coating the droplets greatly decreases the sensitivity while the microdroplets would otherwise sense at these concentrations. Scale bars 25 $\mu$m.

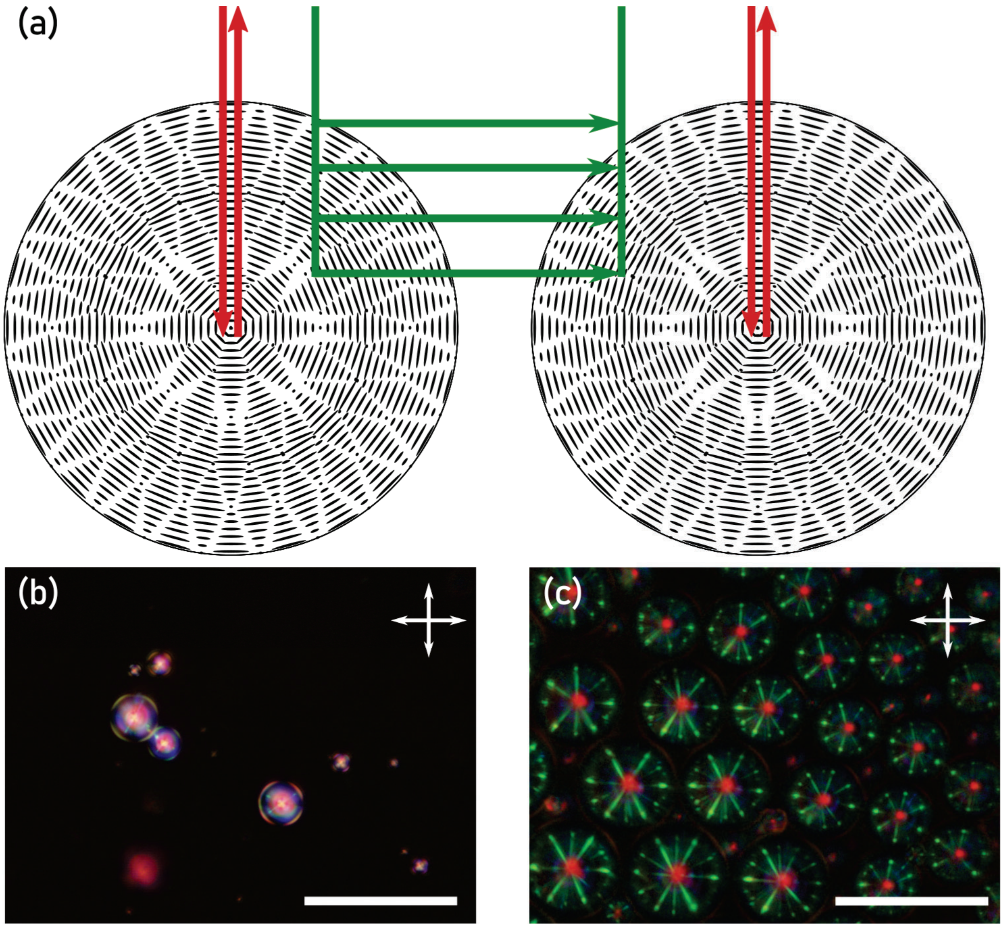


**Figure S4.** Inter-droplet reflections between CLC droplets lead to the unique signatures without adsorbed amphiphile. (a) Light normally incident on the red-reflecting CLC will be reflected as red, while the reflections between droplets will appear green. (b) In a sample with a low density of droplets, the main characteristic of the reflection is single red spots at the droplet center. (c) High droplet densities, however, increases the number of inter-droplet reflections, thus producing a characteristic 'starburst' pattern. Scale bars 50 $\mu$m. (b) and (c) viewed between crossed linear polarizers.

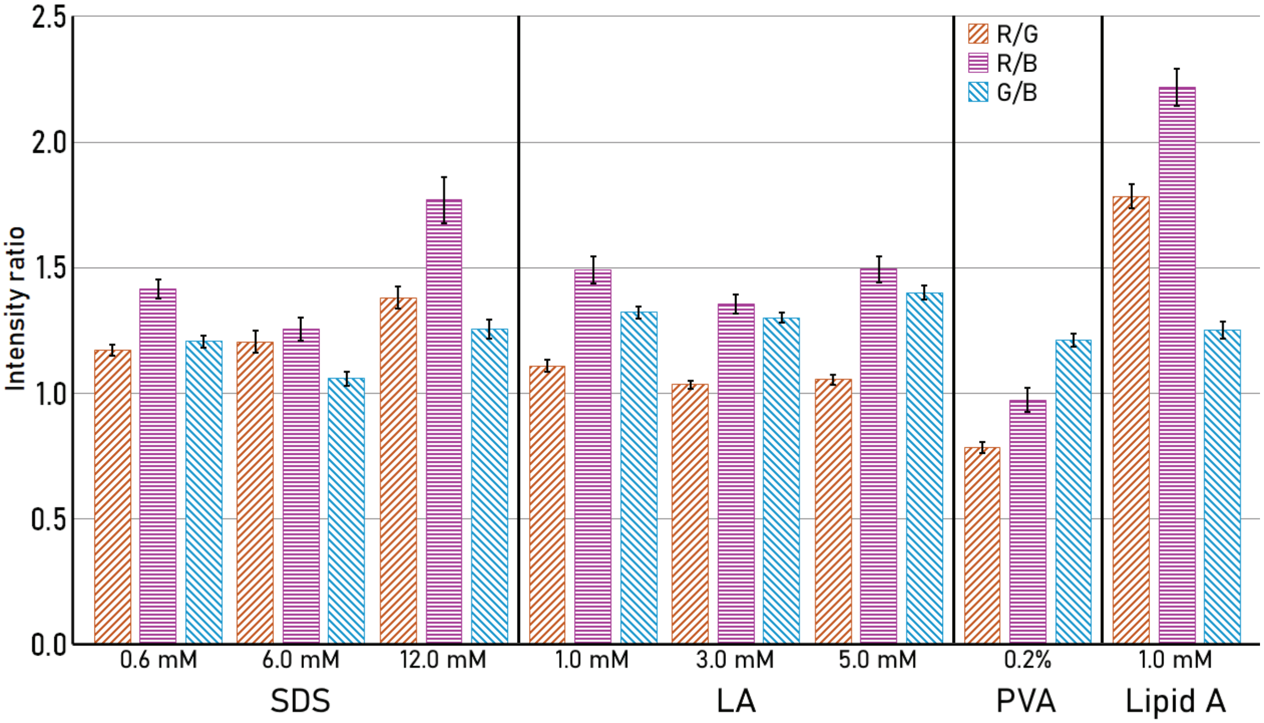


**Figure S5.** Normalized R/G, G/B, and B/G ratios for CLC droplets exposed at varied concentrations of each of the amphiphiles (SDS, lauric acid, Lipid A, and PVA). At least 50 droplets were measured for each group. Error bars indicate the calculated standard error of the mean.

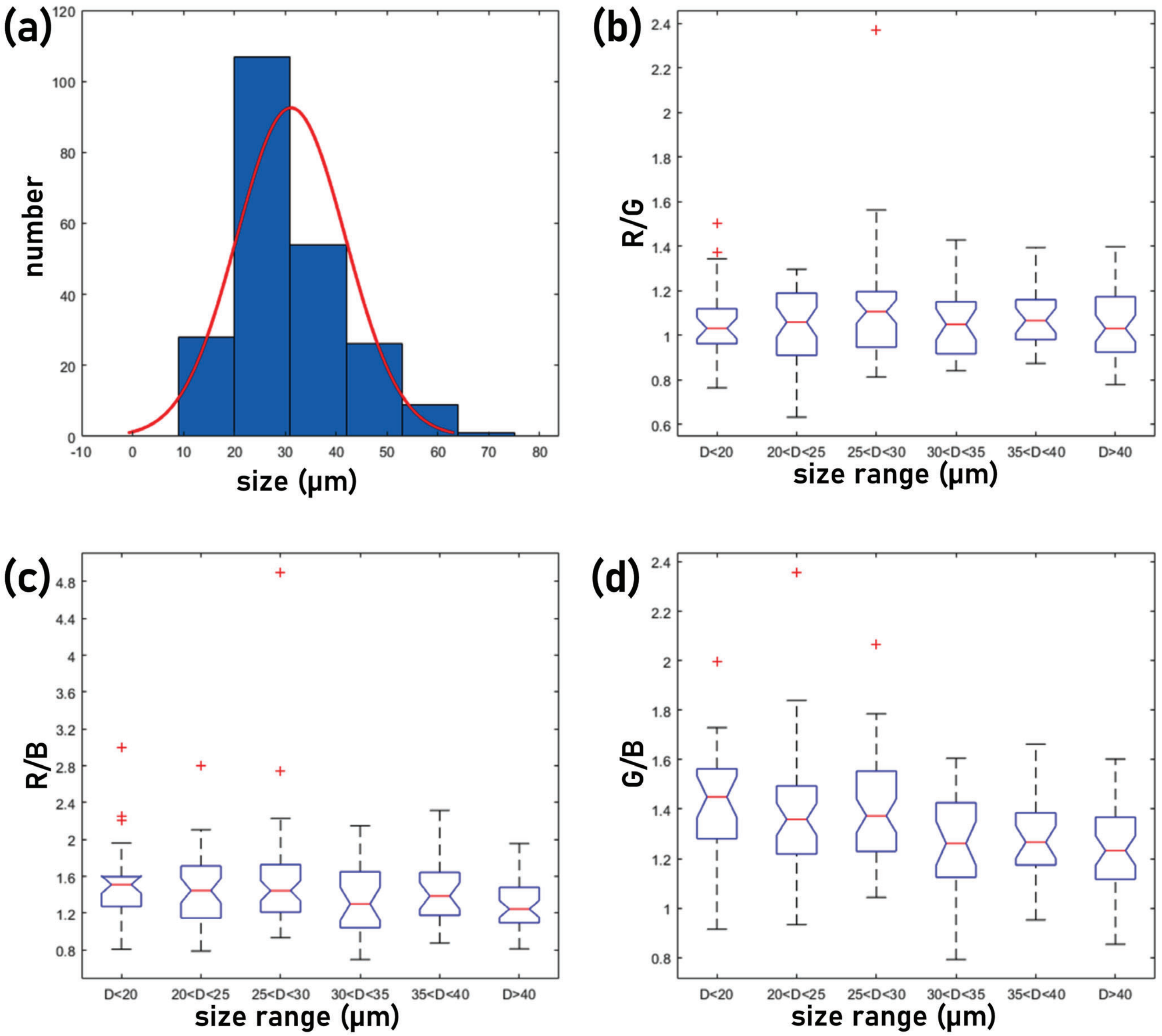


**Figure S6.** (a) Histogram showing the combined size distribution of droplets produced in 1.0, 3.0, and 5.0 mM lauric acid solutions, showing a maximum of droplet distribution between 20-40 $\mu$m. (b-d) Box-and-whisker plots showing the effects of the size distribution on the (b) R/G; (c) R/B; and (d) G/B channel ratios. We saw no effects of size on the channel intensities, showing the final colours were droplet size-independent.

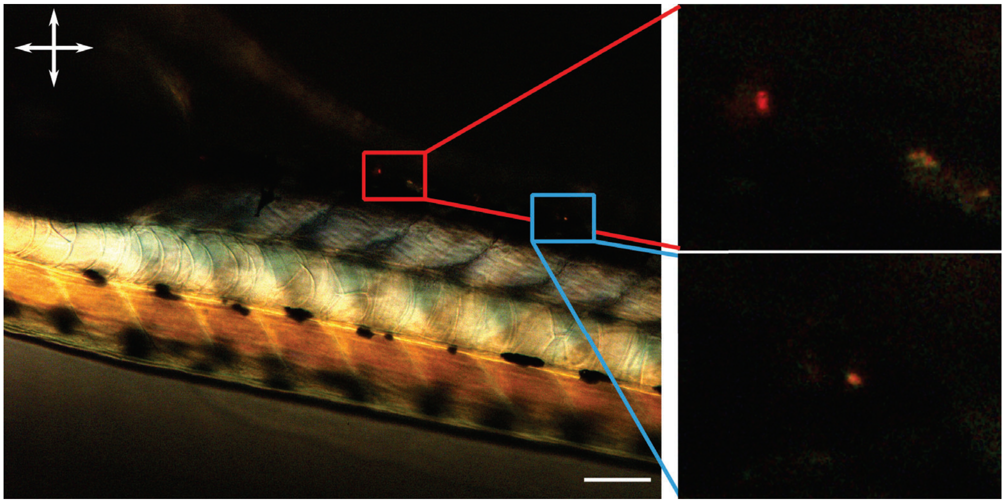

**Figure S7.** A POM micrograph of a zebrafish, viewed between crossed polarizers, showing the birefringence of both the cartilage/bones and the melanocytes (insets). Scale bar 25 $\mu$m.

### Video Captions

**Video S1**: Rehydration of a CLC droplet array, suspended in 0.2% PVA solution and dried on a glass slide, with 0.6 mM SDS solution. Viewed in reflection mode between crossed linear polarizers. Video accelerated 20$\times$.

**Video S2**: Rehydration of a CLC droplet array, suspended in 0.2% PVA solution and dried on a glass slide, with 6.0 mM SDS solution. Viewed in reflection mode between crossed linear polarizers. Video accelerated 5$\times$.

**Video S3**: Rehydration of a CLC droplet array, suspended in 0.2% PVA solution and dried on a glass slide, with 12.0 mM SDS solution. Viewed in reflection mode between crossed linear polarizers. Video accelerated 5$\times$.

**Video S4**: Rehydration of a CLC droplet array, suspended in 0.2% PVA solution and dried on a glass slide, with 5.0 mM lauric acid solution. Viewed in reflection mode between crossed linear polarizers. Video accelerated 5$\times$.
